# Non-additive QTL mapping of lactation traits in 124,000 sequence-imputed cattle reveals novel recessive loci

**DOI:** 10.1101/2021.08.30.457863

**Authors:** Edwardo GM Reynolds, Thomas Lopdell, Yu Wang, Kathryn M Tiplady, Chad S Harland, Thomas JJ Johnson, Catherine Neeley, Katie Carnie, Richard G Sherlock, Christine Couldrey, Stephen R Davis, Bevin L Harris, Richard J Spelman, Dorian J Garrick, Mathew D Littlejohn

## Abstract

Deleterious recessive conditions have primarily been studied in a Mendelian disease context. Recently, several large effect, deleterious recessive mutations were discovered via non-additive GWAS of quantitative growth and developmental traits in cattle. This showed quantitative traits can be used as proxies of genetic disorders if they are indicative of whole animal health status and susceptible to underlying genetic conditions. Lactation traits might also reflect genetic disorders in cattle, given the increased energy demands of lactation and the substantial stresses imposed on the animal. Here, we report a screen of over 124,000 cows for recessive effects based on lactation traits. We discovered novel loci associated with five large recessive impacts on milk yield traits represented by missense variants (DOCK8, IL4R, KIAA0556, and SLC25A4) or premature stop variants (ITGAL, LRCH4, and RBM34) as candidate causal mutations. On milk composition traits, we identified several small effect dominance contributions to previously reported additive QTL. In contrasting analyses of milk yield and milk composition phenotypes, we note differing genetic architectures. Milk yield phenotypes presented lower heritabilities and fewer additive QTL, but higher non-additive genetic variance and a higher proportion of loci exhibiting dominance compared to milk composition phenotypes. Large-effect recessive QTL are segregating at surprisingly high frequencies in cattle. We speculate that the differences in genetic architecture between milk yield and milk composition phenotypes derive from underlying dissimilarities in the cellular and molecular representation of these traits. Lactation yields may act as a better proxy than milk composition traits for a wide range of underlying biological disorders affecting animal fitness

## Background

Non-additive genetic effects are best known and studied in Mendelian disease contexts, where recessive conditions have been shown to have major deleterious impacts on the health and performance of animals. These studies have mostly used a ‘forward genetics’ approach, where observation of a disease phenotype precedes fine mapping and sequencing to highlight the mutation [1–3]. The reverse approach has also been applied, where candidate loss of function genotypes were identified and phenotyping was subsequently conducted to detect the impact of the mutation [4,5]. Though examples remain limited, genome-wide association approaches have been used to investigate non-additive effects in quantitative traits. Recent studies include the investigation of complex traits in both humans [6] and cattle [7–11]. Reynolds *et al*. identified several recessive mutations in cattle with major negative impacts on growth and developmental traits, where some of these loci represented underlying genetic disorders [11].

The concept of using routinely gathered, quantitative traits as proxies of genetic disorders is based on the idea that phenotypes such as growth or liveweight can be indicative of whole animal health status, where reduced growth might be due to some underlying genetic disorder, and that those effects could be detected via GWAS. It is therefore of interest to consider what other traits might serve as proxies of animal fitness, with a view to extend the utility of this approach. Lactation traits such as milk volume comprise one of the most commonly targeted classes of quantitative traits studied in cattle, where additive analyses of these traits have presented numerous candidate causative genes such as *DGAT1* [12], *GHR* [13], *ABCG2* [14], *GPAT4* [15], and *MGST1* [16]. Lactation traits might also be reflective of genetic disorders, given the increased energy demands of lactation and the substantial metabolic and physiological stresses imposed on the animal [17]. We wondered therefore whether the application of non-additive models to lactation data might identify further recessive mutations, and to this end, have conducted non-additive GWAS for milk traits in 124,000 animals. We contrast the additive and non-additive genetic architectures of milk yield traits and milk composition traits. Finally, we describe the discovery of novel major effect recessive loci, highlighting candidate mutations that potentially underlie undiagnosed recessive disorders.

## Methods

### Animal populations

The dataset reported in this study consists of 124,364 New Zealand dairy cattle. These animals come from a mixed breed population, where 20,893 are 16/16^th’^s Holstein-Friesian (HF), 13,184 are 16/16^th’^s Jersey (J), 67,520 are crosses involving varying proportions of the two breeds (HFXJ), and 22,767 are HF or J crossbreeds with minor proportions of other breeds including Ayrshire, Brown Swiss, or Hereford (and other crosses). An individual’s breed may be coded as 16/16^th^s, however, this does not preclude the possibility that an ancestor may be crossbred as matings between 15/16^th^s and 16/16^th^s animals result in 16/16^th^s offspring. The animals were born between 1990 and 2018 with a mean birth year of 2010.

### Phenotypes

Five first-lactation milk phenotypes were investigated in this study. These include three milk yield traits; milk volume (L/Lactation; a lactation refers to a standardised 268 day lactation; N = 124,356), milk protein yield (kg/Lactation; N = 124,356), and milk fat yield (kg/Lactation; N = 124,356), and two milk composition traits; milk protein percentage (%; N = 124,363), and milk fat percentage (%; N = 124,363). Milk protein yield and milk fat yield are the product of the milk volume multiplied by the milk protein percentage or milk fat percentage, respectively.

Prior to genetic analysis, phenotypes were adjusted based on effects obtained from the national genetic evaluation of the entire cattle population (30 million animals) which fits mixed linear models. Fixed effects in that model included contemporary group, age at calving, stage of lactation, and record type (records may be made at am milkings, pm milkings, or both). Since animals have varying numbers of herd-test measurements within each milk trait, these were aggregated to a phenotypic deviation such that each animal has a single record and a corresponding weighting reflecting the amount of information in the record [18].

### Sequence-based imputation reference panel

Whole genome sequencing was performed on 1,300 animals that were mostly ancestral sires, these animals comprised the reference population for sequence-based imputation. Animals comprising HF (N=306), J (N=219), HFXJ (N=717), or other breeds and crossbreeds (N = 58) were sequenced on Illumina HiSeq 2000 instruments targeting 100bp paired-end reads. Sequence data were aligned to the ARS-UCD1.2 reference genome assembly using BWA 0.7.17 [19] resulting in a mean read depth of 15x. Variant calling was performed using GATK v4.0.6.0 [20], followed by variant filtering via Variant Quality Score Recalibration. Using animals with high read depth (>10x, N = 850), variants were filtered out if they were singletons, were multi-allelic, had a map quality score lower than 50, or had a Mendelian error rate above 5%. These criteria left 21,005,869 whole genome sequence variants from the 850 highest read depth animals, where these positions were then extracted from the sequence data on all 1,300 animals and phased using Beagle 5.0 [21] to create the sequence-based imputation reference panel.

### Genotyping

The study animals (N = 124,364) were genotyped using SNP chips, where either ear-punch tissue samples or blood samples were used for DNA extraction. Genotyping was performed using a variety of platforms including GeneSeek GGPv1, GGPv2, GGPv2.1, GGPv3, GGPv3.1, GGPv4, GGP50kv1, GGP50kv1.1, Illumina BovineSNP50v1, Illumina BovineSNP50v2, or BovineHD SNP-chips. Samples were processed for DNA extraction at GeneMark (Hamilton, New Zealand) using Qiagen BioSprint kits or GeneSeek (Lincoln, NE, USA) using Life Technologies’ MagMAX system.

### Consolidation of SNP-chip panels for sequence imputation

Imputation from genotyping panels to sequence resolution was performed as described in Wang *et al*. [22]. Genotype panels were grouped into four sets; GGP panels (GGPv1, GGPv2, GGPv2.1, GGPv3, GGPv3.1, and GGPv4), 50K panels (BovineSNP50v1, and BovineSNP50v2), GGP50k panels (GGP50kv1, GGP50kv1.1), and the BovineHD panel. Animals genotyped on the GGP panels were imputed to the BovineSNP50v1 panel, then combined with the physically genotyped 50K panel animals and further imputed to the BovineHD panel. Animals genotyped on the GGP50k panels were separately imputed to the BovineHD panel. In order to incorporate the large amount of custom content genotyped on the GGPv3 platform, we conducted similar imputation steps to impute all animals to GGPv3. We then combined the imputed and physically genotyped panels (imputed HD, imputed GGPv3, and physically genotyped HD), and imputed these animals to sequence resolution using the sequence-based imputation reference population, described above. Post-imputation filtering to remove very rare variants (homozygous alternate count ≤ 5) was performed, as well as a filter to remove variants that imputed poorly based on the dosage R^2^ statistic (DR^2^; DR^2^ < 0.7). After the application of these filters, 16,640,294 variants remained for GWAS and further analysis.

### Genotypes for population structure adjustment

We used content from the Bovine SNP50 chip platform to account for the population structure of the sample. From the initial 54,708 autosomal SNPs, we filtered to remove markers with high missing genotype rates (> 0.01), low minor allele frequency (< 0.02), or high deviations from expected Hardy-Weinberg equilibrium (> 0.15, calculated within breed). This was followed by further filtering to remove markers that appeared to impute poorly (DR^2^ > 0.9), and markers in high LD with another marker on the panel (pairwise R^2^ > 0.9, within 1 Mbp). These criteria resulted in a set of 31,451 SNP chip markers for subsequent analysis.

### Heritability estimates

We estimated breed-specific additive and dominance heritabilities using genomic relationship matrices (GRMs) using GCTA software [6,23]. Variance components were estimated from purebred individuals (HF = 20,893, J = 13,184), using the same set of 31,451 filtered BovineSNP50 SNPs used for population structure adjustment (filters described in the previous section). GCTA estimates variance components using a restricted maximum likelihood (REML) approach, where additive heritability (h^2^) is the ratio of additive genetic variance to phenotypic variance, and dominance heritability (δ^2^) is calculated as the ratio of dominance genetic variance to phenotypic variance.

### GWAS

#### Model Overview

We applied a non-additive GWAS method similar to that described in Reynolds *et al*. [11] to identify non-additive QTL for milk traits. This two-step method first uses a leave-one-segment-out (LOSO) approach to fit genomic marker effects to adjust for population structure, and a second-step Markov chain Monte Carlo (MCMC) method to test the effects of all imputed-to-sequence variants, one at a time. In general, for each sequence variant the method fits the following model:

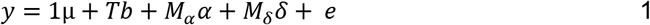

Where ***y*** indicates a vector of one of the 5 phenotypes of interest, pre-adjusted as described in the ‘*Phenotypes*’ section above, ***μ*** is the overall mean, ***1*** is a vector of ones, ***b*** is a vector of genotype class effects for the sequence variant of interest, and ***T*** is the design matrix relating records to genotype class for the sequence variant. The vector ***α*** represents random SNP chip additive marker effects spanning the whole genome except the segment of interest such that ***α*** ~ N(***0***, ***I***σ_α_^2^), where ***I*** is an identity matrix of order equal to the number of marker effects and σ_α_^2^ represents the additive marker effect variance, ***δ*** is a vector of random SNP chip dominance marker effects spanning the whole genome except the segment of interest such that ***δ*** ~ N(***0***, ***I***σ_δ_^2^), where σ_δ_^2^ represents the dominance marker effect variance. ***M_α_*** and ***M_δ_*** are matrices with each column representing the covariate values for a marker locus ([0, 1, 2] and [0, 1, 0], respectively). The vector ***e*** represents residuals with ***e*** ~ N(***0***, ***R***), where for a simple model based on single observations ***R*** = ***I***σ_e_^2^, where ***I*** is an identity matrix of order equal to the number of phenotypic records and σ_e_^2^ represents the residual error variance. Since the traits investigated here are represented by the mean of a variable number of repeated observations, the diagonal elements of ***R*** varied according to the number of observations contributing to the yield deviation. One notable contrast to the model implemented in Reynolds *et al*., is that in the current model, we fit both additive (M_α_) and dominance (M_δ_) effects of the genomic markers to adjust for population structure. This modification was made to better control the inflation observed when analysing milk traits in a population larger than that studied in Reynolds *et al*. [11].

##### Population structure adjustment

500 samples of vectors of plausible marker effects, ***ᾶ*** and 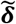, for the 31,451 SNP-chip markers, were generated using single-site Gibbs sampling from an extension of the BayesC0 algorithm implemented in GenSel using standard priors [24]. That algorithm was performed while omitting the ***Tb*** term from (1) and convergence of the Markov chain of plausible marker effects was determined using the Geweke diagnostic [25]. LOSO was used to avoid fitting SNP-chip marker effects in linkage disequilibrium with the sequence variant being tested. The genome was partitioned into 10Mbp LOSO intervals and, for each interval, phenotypes were adjusted for the samples of SNP chip marker effects except those within the relevant LOSO interval. This produced distinct LOSO-adjusted phenotypic deviations for each 10Mbp interval for each sample of plausible marker effects.

##### Association analysis

For each sequence variant, we sampled genotype class effects for each plausible sample of LOSO-adjusted phenotypic deviations. We obtained MCMC chains of additive and dominance genotypic effects, and standard-additive effects as contrasts of these plausible genotype class effects. These posterior distributions were summarised by their posterior means, posterior standard deviations, and z-statistics following a standard Normal distribution [26]. The statistical significance of standard-additive, additive, and dominance genetic effects were evaluated using a Z-test.

### QTL identification, significance criteria, and annotation

We primarily aimed to detect non-additive QTL, as such we declared variants significant if the dominance genotypic effect, **d**, passed a false discovery rate (FDR) threshold of 1 × 10^−3^. For each phenotype, this FDR threshold was calculated using q-values [27] as implemented in the *qvalue* package in R [28]. Since we were particularly interested in medium- to large-effect QTL, only loci with effect sizes (**a**, or **d**) greater than 5% the phenotypic standard deviation of the trait were considered for further downstream analyses. We calculated the dominance coefficient 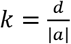 each significant QTL to characterise the non-additive mechanism presented, where *k* ≈ 0 represents a completely additive locus, *k* ≈ 1 represents a completely recessive locus, *k* < 1 a partially dominant locus, and *k* > 1 an over-dominant locus.

For standard-additive effects, **α**, we used GCTA-COJO [29] to detect tag variants for QTL identified in our standard-additive GWAS. GCTA-COJO utilises LD structure and GWAS summary statistics to iteratively identify significant QTL at the FDR threshold of 1×10^−3^. We used sequence annotations from variant effect predictor (Ensembl 97, [30]) to highlight mutations that might be responsible for non-additive QTL identified, where the potential impact of missense mutations on protein function was judged using SIFT scores [31].

### Iterative GWAS

We aimed to investigate whether multiple dominance QTL might segregate at associated loci, so implemented an iterative GWAS approach to differentiate QTL. Here, we first identified the variants on each chromosome that surpassed the false discovery threshold. We then adjusted the phenotype for the genotype class effects of the most significant variant (or candidate causal variant if identified) and then re-ran the GWAS model on the chromosome of interest using the residual phenotype. This process was iterated until there were no further significant QTL on the chromosome.

## Results

### Heritabilities of lactation traits

We first estimated additive and dominance heritabilities for each phenotype within each breed to investigate the additive and non-additive genetic architecture of each trait. These results are shown in Table 1, additive heritabilities far outweighed dominance heritabilities, though presented ratios of similar magnitude to those previously reported for other traits and populations [8,32]. Milk fat yield in Jersey cows had the highest dominance heritability at 0.074, and milk protein percentage in Holstein-Friesian cows had the lowest dominance heritability at 0. Of note, a distinct contrast in relative heritabilities was apparent between milk composition and milk yield traits, where composition traits had high additive heritabilities but near zero dominance heritabilities, and yield traits presented lower additive heritabilities but higher dominance heritabilities (Table 1).

**Table 1.** Heritability estimates for lactation traits

| Table 1 \| Heritability estimates for lactation traits |  |  |  |  |
| --- | --- | --- | --- | --- |
| Trait | $h^2_{HF}$ | $\delta^2_{HF}$ | $h^2_J$ | $\delta^2_J$ |
| Milk Volume | 0.296 | 0.044 | 0.312 | 0.064 |
| Milk-Fat Yield | 0.261 | 0.059 | 0.232 | 0.074 |
| Milk-Protein Yield | 0.235 | 0.053 | 0.236 | 0.073 |
| Milk-Fat Percentage | 0.7 | 0.006 | 0.616 | 0.015 |
| Milk-Protein Percentage | 0.642 | 0 | 0.636 | 0.005 |

$h^2$ - Additive heritability, $\delta^2$ - dominance heritability, HF - Holstein-Friesian, J - Jersey

### Lactation trait GWAS

We performed GWAS’ across the five milk traits of interest, namely milk volume, milk protein yield, milk fat yield, milk protein percentage, and milk fat percentage to identify non-additive QTL (Figure 1). Both additive and dominance effects are included in these plots, where iterative analysis identified 23 dominance QTL signals that passed our FDR threshold. These included 10, 11, 12, 8, and 7 QTL from 4,618, 2,706, 8,525, 8,987, and 5,800 significant variants across milk volume, milk protein yield, milk fat yield, milk protein percentage, and milk fat percentage, respectively. These signals spanned 13 discrete autosomes. In standard-additive GWAS, following iterative COJO analysis, we identified 217, 152, 142, 673, and 457 QTL across milk volume, milk protein yield, milk fat yield, milk protein percentage, and milk fat percentage, respectively.

**Figure 1.**
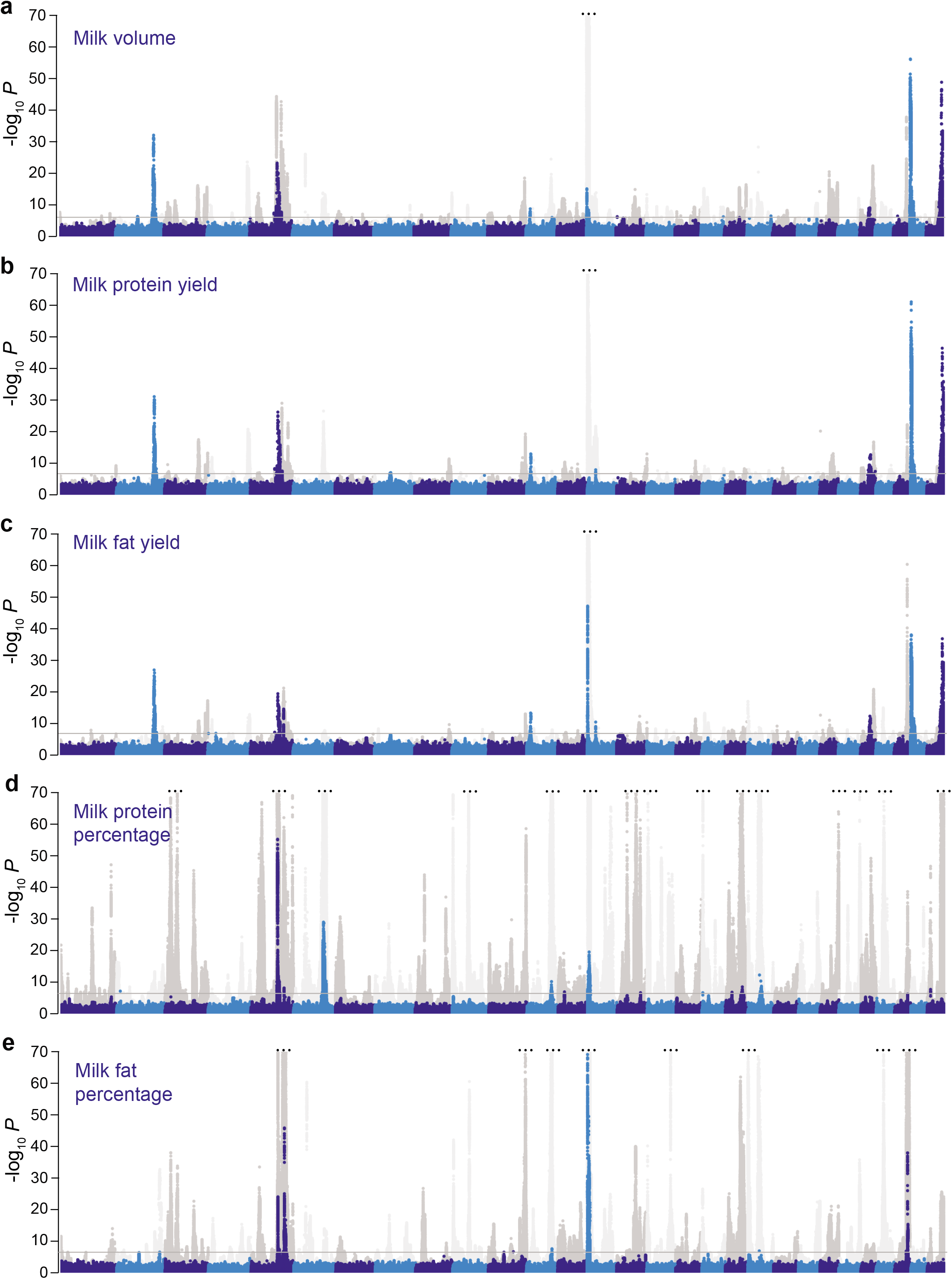
Dominance and additive Manhattan plots for lactation traits. a-e, Manhattan plots for milk volume (a), milk protein yield (b), milk fat yield (c), milk protein percentage (d), and milk fat percentage (e) showing significance of genotypic dominance (blue and light blue), and additive (grey and light grey) estimates for ~16.6 million imputed sequence variants. Chromosomes are differentiated by alternating colours and a grey line indicates the false discovery rate of 1×10^−3^, used to account for multiple testing. The y-axes are truncated for display purposes (indicated by 3 dots).

### Dominance QTL

We identified 15 significant dominance QTL for milk yield traits, and 11 for milk composition traits (Table 2, Supplementary Table 1). Across the milk yield dominance QTL, the majority (N=12) were recessive effects and they were located on chromosomes 2, 4, 5, 8, 12, 25, 28, and 29. Seven of these signals appear to be novel to the current study, the remainder having been recently highlighted in our analysis [11] of growth and developmental traits in an overlapping population to that described here. Across the 11 milk composition dominance QTL, the majority (N=8) presented partial dominance effects, with six of these representing loci identified from previously published additive GWAS (Supplementary Table 1).

**Table 2.**
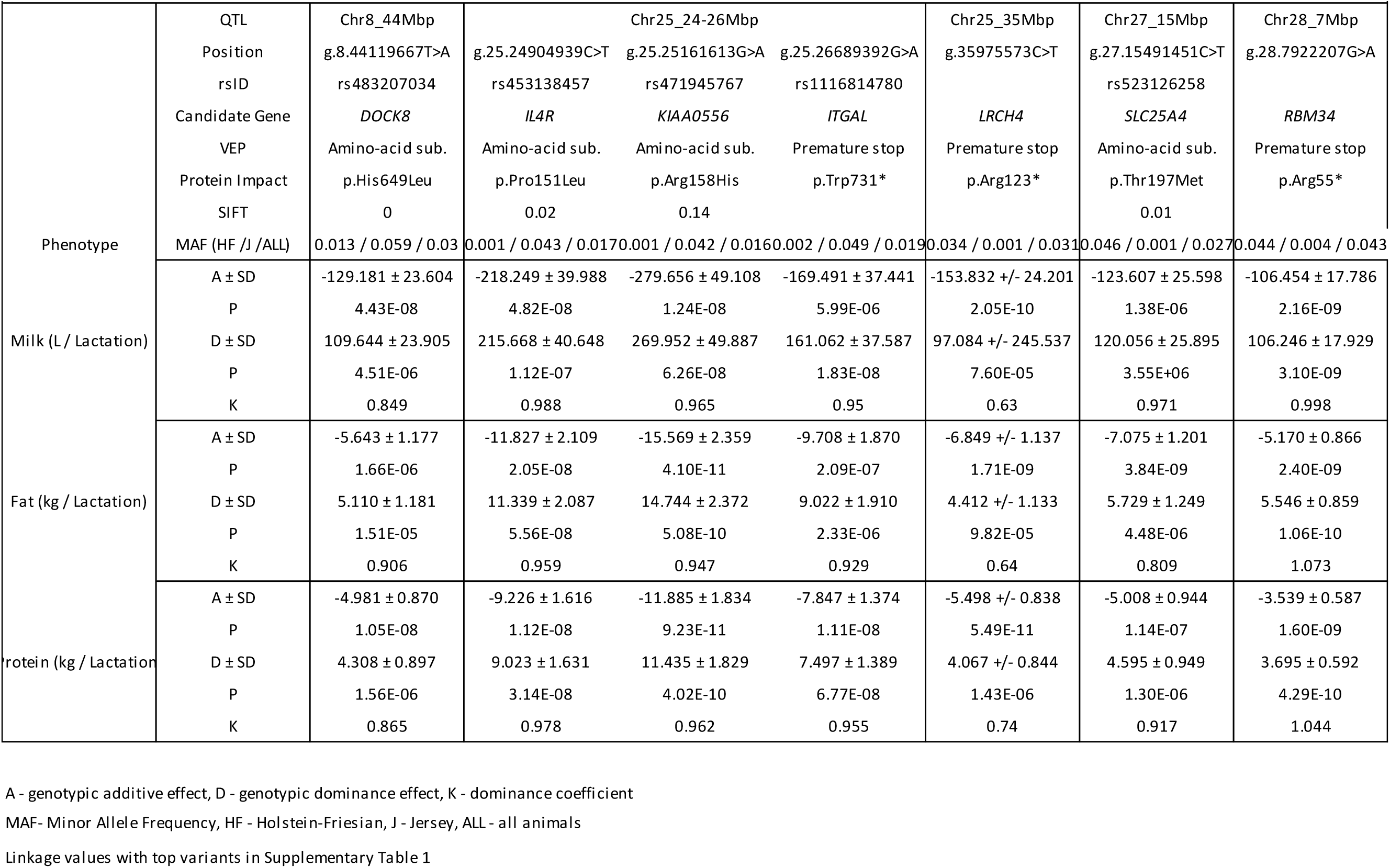
Association statistics for candidate mutations at recessive loci

Figure 2a contrasts the minor allele frequency and effect size of dominance components for all of these effects. Interestingly, milk composition trait QTL appeared to be tagged by high minor allele frequency variants with comparatively small effect sizes, whereas milk yield QTL tag variants had low minor allele frequencies and larger effects. The type of effects also appeared to differ between traits (Figure 2b), where we noted an abundance of recessive QTL in milk yield traits, whereas milk composition traits mostly comprised partially dominant QTL.

**Figure 2.**
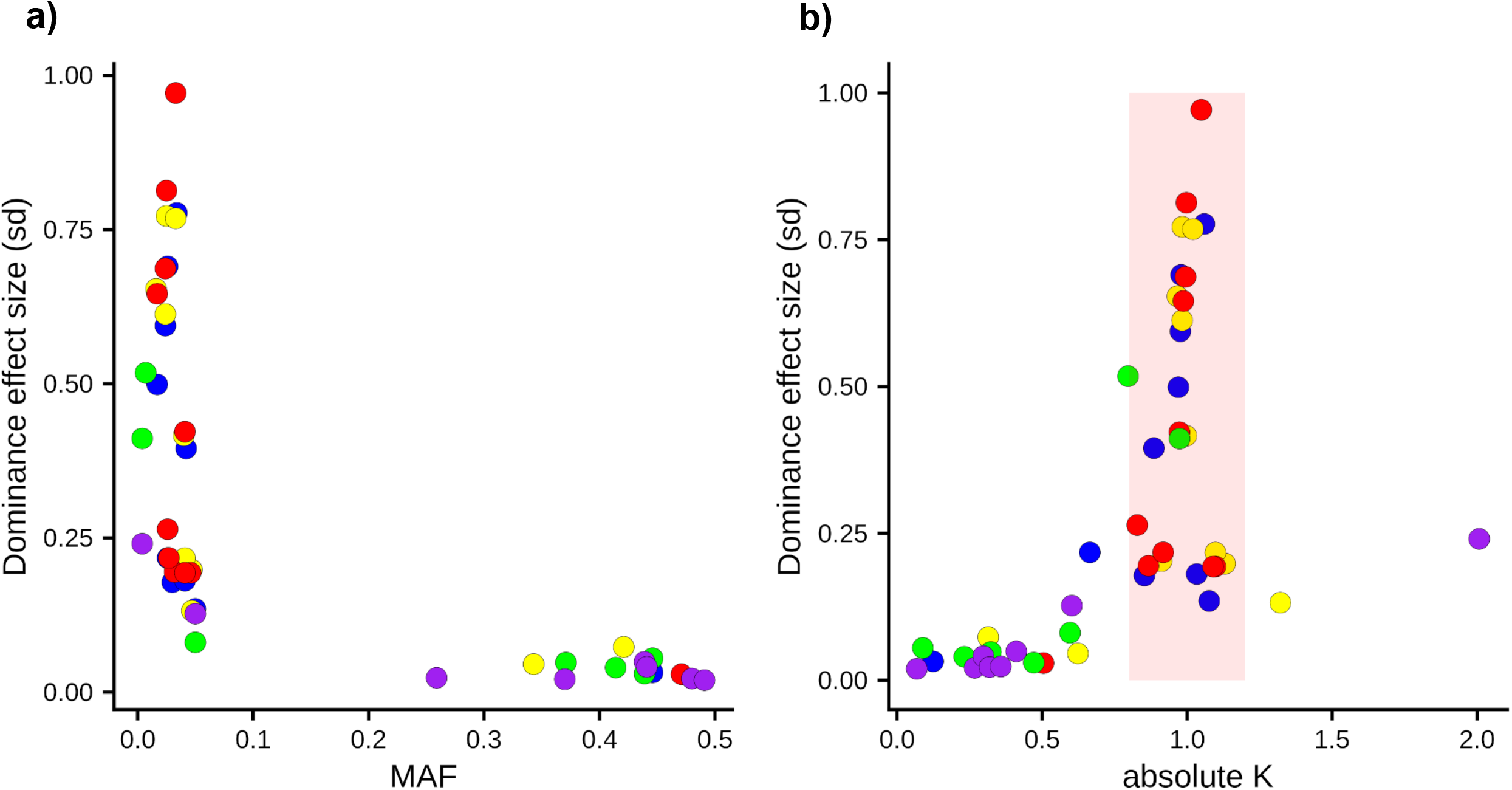
Plots presenting genetic architecture of significant dominance QTL from GWAS on milk volume (milk), milk protein yield (prot), milk fat yield (fat), milk protein percentage (protper), milk fat percentage (fatper), and. The plots contrast the minor allele frequency (MAF) against the dominance effect size (a), and the absolute value of k, where k = d/|a| against the dominance effect size (b).

### Candidate causal mutation identification

Given the status of recessive milk yield QTL as potentially representing novel bovine disorders, we prioritised these QTL for further investigation, selecting QTL where the dominance coefficient (*k*) was near 1 (0.7 < *k* < 1.3). We used sequence annotations from variant effect predictor to highlight mutations that might be responsible for these effects (Ensembl 97, [30]), highlighting variants that were in strong to moderate LD (R^2^ > 0.7) with the lead variant per locus, and that were also predicted to alter or disrupt protein function. We identified 5 novel recessive QTL (including one biologically compelling near-significant recessive QTL), and several other recessive QTL previously identified and attributed to mutations in the *PLCD4, FGD4, MTRF1, GALNT2, DPF2*, and *MUS81* genes [11]. Figure 3 presents the position, regional LD, and association statistics for the QTL novel to the current study. Note that we have applied relatively simple annotation criteria and only highlight protein-coding variants as candidates since, for recessive signals at least, we consider protein altering mutations primary candidates given the loss of function connotation for these effects. Supplementary Table 1 shows all significant QTL identified, including those not expanded upon here.

**Figure 3.**
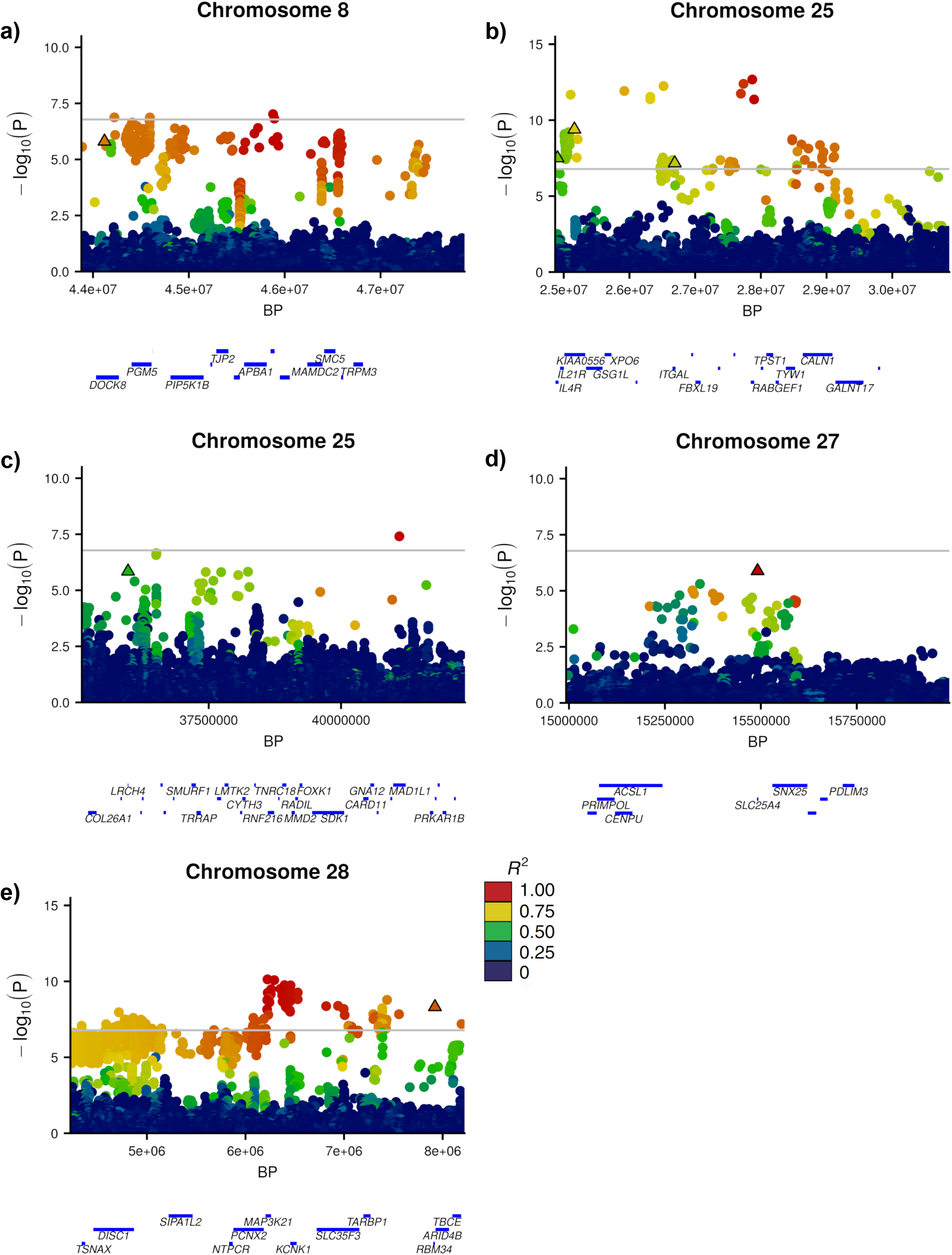
Manhattan plots for the five novel milk protein yield QTL representing the chr8:44Mbp (a), chr25:24-27Mbp (b), chr25:35Mbp (c), chr27:15Mbp (d), and chr28:7Mbp (e) loci. Variants are coloured by LD (R^2^) values with the top tag variant per locus, protein coding variants are shown as outlined triangles. Gene tracks are presented below each plot based on Ensembl 97, where gene names have been filtered on size.

### Chromosome 8

Chromosome 8 presented a significant signal at 45Mbp for milk protein yield and milk fat yield. The most significant variants for these signals (g.45878531A>C and g.45880948C>T) were in strong LD (R^2^=0.99), and we note an annotated missense variant (g.44119667T>A, rs483207034) in high LD with both top-associated variants (R^2^ = 0.85 and 0.85, respectively; Figure 3a). This variant in the *DOCK8* gene results in an amino acid (p.His649Leu) change and has a predicted deleterious impact (SIFT = 0).

### Chromosome 25

A dispersed QTL signal is apparent on chromosome 25 at 24-27Mbp across the three lactation yield traits, with the top variants at g.25921991AT>T for milk fat yield, and g.27868969C>T for milk protein yield and milk volume. Effect prediction highlighted three candidate causal mutations in the region. These included a p.Pro151Leu substitution in the *IL4R* gene (g.24904939C>T, rs453138457) with R^2^ = 0.74, and 0.62, for the milk fat and milk protein/milk volume top variants, respectively, another missense variant (p.Arg158His) in the *KIAA0556* gene (g.25161613G>A, rs471945767) with R^2^ = 0.89, and 0.74, respectively, and a nonsense variant (p.Trp731*) in the *ITGAL* gene (g.26689392G>A, rs1116814780) with R^2^ = 0.76, and 0.70, respectively (Figure 3b). While these are plausible candidates to explain the QTL, we were not able to distinguish between the candidates through iterative analysis, where fitting any one of these candidates removed the majority of the association at this locus.

A second signal for protein yield on chromosome 25 was observed at 35Mbp. This locus maintained its significance after accounting for the chromosome 25 25Mbp QTL through iterative analysis, suggesting it was a discrete effect. The locus presented a strong candidate causative mutation as potentially underlying the effect, comprising a stop gain mutation (g.35975573C>T; Arg123*) in the *LRCH4* gene that was the third most highly associated variant at this locus overall (Figure 3c). We observed a mostly recessive effect for this variant (k = 0.74), where animals carrying the heterozygote and homozygous alternate genotypes produce 1.44kg, and 11.21kg less milk protein per lactation compared to the homozygous reference genotype. When fitting g.35975573C>T as a fixed effect, the significance of the QTL is removed, and no further QTL are apparent on the chromosome (Supplementary Figure 1).

### Chromosome 27

We observed a signal at 15Mbp on chromosome 27 for milk protein yield. Although this did not surpass our q-value FDR threshold of 1×10-3 (equivalent to P = 1.65×10^−7^), this signal was conspicuous given that the lead variant (g.15491451C>T; rs523126258, p-value = 1.30×10^−6^) is a predicted deleterious missense mutation (p.Thr197Met) in the *SLC25A4* gene. Figure 3d shows a Manhattan plot for this region.

### Chromosome 28

We previously reported a major recessive bodyweight QTL on Chromosome 28 represented by a likely causative splice acceptor mutation in *GALNT2* (g.2281801G>A) [11]. This QTL was apparent in the current analysis, impacting all three milk yield traits. However, the application of iterative association analysis revealed a secondary QTL approximately 4Mb downstream of the *GALNT2* mutation at Chr28:6-7Mbp (top variant at g.6223350G>A). This residual signal highlighted a stop-gain non-sense mutation (g.7922207G>A) strongly linked to the g.6223350G>A variant (R^2^ = 0.89; Figure 3e). This stop-gain mutation (p.Arg55*) is in the *RBM34* gene, and appears to be in linkage equilibrium with the *GALNT2* causal mutation (R^2^ < 0.001), having little association with bodyweight in our previous analysis (p=0.37;[11]). Upon the second chromosome 28 GWAS iteration (fitting both *GALNT2* and *RBM34* mutations as fixed effects), there were no further significant QTL on the chromosome (Supplementary Figure 2).

### Dominance QTL for composition traits

In addition to the recessive QTL identified for milk yield traits, we also identified dominance QTL for milk composition traits. We investigated these effects and observed several partial dominance QTL in close proximity to previously described additive loci. The tag variants of these QTL were adjacent the genes; *CSF2RB* [33], *MGST1* [16], *DGAT1* [12], *GHR* [13], *GPAT4* [15], and *PICALM* [34] and, in each case, these variants were in high linkage disequilibrium (R^2^ > 0.8) with previously identified causal and/or tag variants (Supplementary Table 1).

Milk protein percentage presented multiple dominance QTL on Chromosome 6 within the Chr6:80-85Mbp region (Supplementary Table 1). The most significant of these QTL presented the top variant g.84112451C>A and shows a partial dominance effect. Unlike the examples highlighted above, no very strongly linked candidate mutation was identified, though we note that this variant is in moderate LD with a previously proposed causative variant in *CSN1S1* (R^2^ = 0.53; p.Glu192Gly mutation; g.85427427A>G) [35]. Chromosome 12 presented a significant dominance QTL, where we observed a partial dominance effect at 68Mbp for milk protein percentage with the top variant at g.68763031T>TG. As with the chromosome 6 locus, no particularly obvious candidate causal variant or gene was identified that might account for this signal.

### Contrasting additive and dominance GWAS results

Figure 4 compares minor allele frequency (MAF) and the effect sizes between homozygous genotypes across all traits and genetic mechanisms. As might be expected, we observed many more additive QTL than dominance QTL across all traits. Notably however, mutations detected via dominance GWAS in milk yield traits presented very large effects compared to the additive QTL detected for these traits, and most presented a recessive mechanism. On the other hand, the largest effects presented for the two milk composition traits were mostly additive QTL, where dominance effects tended to be higher MAF and incompletely dominant in their presentation of effect.

**Figure 4.**
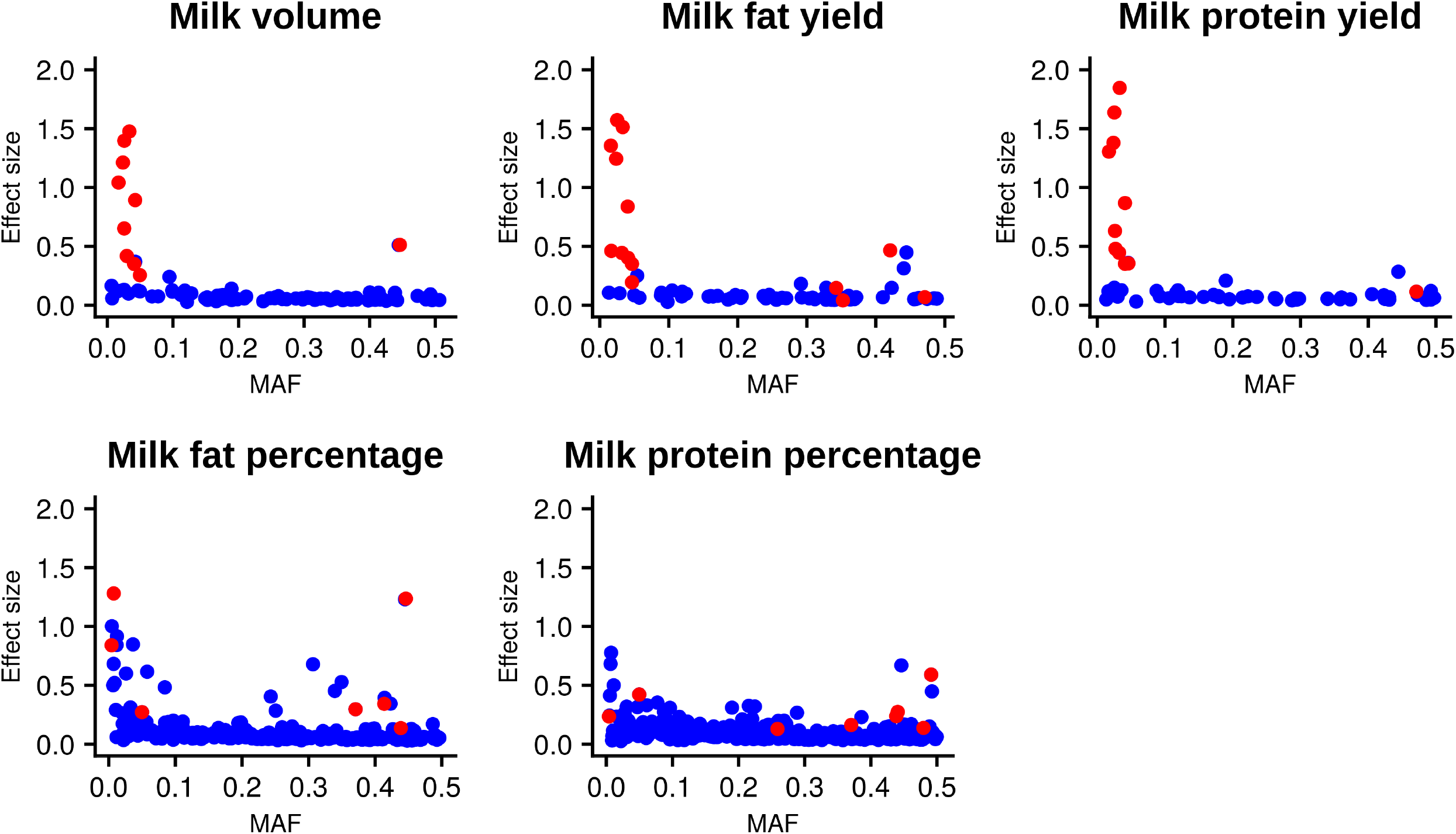
Plots contrasting minor allele frequency (MAF) and the absolute effect size between homozygote genotype classes (Effect size) for additive (blue) and dominance (red) QTL detected via GWAS across lactation traits.

## Discussion

The results in this study highlight the presence of many non-additive QTL for milk traits in cattle. The majority of these signals for milk yield traits present recessive QTL, identifying five novel loci and several previously described recessive QTL [11]. Although milk protein percentage and milk fat percentage traits also yielded many dominance GWAS signals, most presented partially dominant QTL that appeared to represent minor dominance components to previously reported additive QTL.

### Different trait classes present contrasting additive and non-additive genetic architectures

One remarkable observation from the current study is the apparent difference in additive and non-additive genetic architectures between milk yield traits and milk composition traits. Dominance heritabilities of yield traits ranged from 3% to 7%, whereas composition traits have dominance heritabilities at or near zero. By contrast, additive heritabilities ranged from 23% to 31% for yield traits, compared to composition traits which ranged from 64% to 70%. These findings are consistent with Sun *et al*. [8] where they observed similar additive and dominance heritabilities and suggest dominance, in particular recessive mechanisms, may play a bigger role in the regulation of yield traits than composition traits.

These architecture contrasts were also apparent when comparing the properties of individual dominance QTL between milk yield and milk composition traits. Dominance QTL identified in milk yield traits manifested primarily with recessive genetic mechanisms, while milk composition traits presented primarily partial dominance effects. Of further note, milk yield trait dominance QTL typically had low minor allele frequencies and large effect sizes, whereas dominance QTL for milk composition traits were typically characterised by high minor allele frequencies and smaller effect sizes. We theorise that these observations may be due to the way in which the different traits are able to reflect underlying deleterious recessive syndromes – i.e., their utility to serve as proxies of genetic disorders. Of all recessive QTL detected in the current study, we previously validated a subset of these as representing new genetic disorders [11]. Although we did not investigate the novel recessive loci in this study with the same rigour as those investigated in Reynolds *et al*., their very large, uniformly negative effects suggest some at least will similarly validate as new recessive syndromes. Notably, none of these loci (new or old) show substantial effects on milk composition, suggesting milk fat and protein percentage traits do not readily reflect recessive effects. This finding can be rationalised by the comparatively broad range of biological processes reflected by milk yield traits (or the growth and development traits investigated in Reynolds *et al*. 2021), where the energy demands of lactation (or growth) might be expected to manifest a wide range of other organismal stresses. The relative composition of milk components, by contrast, likely represents a narrower spectrum of mammary-specific biology that we hypothesise is less able to serve as a proxy of animal fitness.

It should be acknowledged that given protein yield and fat yield are the products of milk volume and their respective percentages, these traits are not independent. We observe the variance components and genetic architectures of milk fat yield and milk protein yield are more comparable to milk volume than their respective composition traits. This suggests milk volume has a greater influence on milk fat yield and milk protein yield due to the additional environmental factors and measurement errors affecting milk volume.

### Previous studies highlighting recessive effects on quantitative traits

As discussed above, we recently reported an investigation of growth and developmental traits that identified non-additive QTL using similar approaches to those presented here [11]. That study demonstrated how quantitative traits can be used as proxies to map genetic disorders without prior disease identification. In doing so, using sequence-resolution variants, the research highlighted several recessive QTL represented by variants in the *PLCD4, FGD4, MTRF1, GALNT2, DPF2*, and *MUS81* genes, each with large effects on bodyweight and other quantitative traits. The work presented here builds on those findings; we identified many of the same recessive mutations as well as several additional recessive QTL. The additional discoveries made here can be assumed to reflect the increased sample size leveraged in the current study.

Aside from the Reynolds *et al*. study discussed above, few other studies have highlighted major effect recessive impacts using quantitative trait data. Although non-additive GWAS with large sample sizes has been performed in cattle [10,32], low marker densities in these earlier studies may have hampered the ability to directly resolve candidate causative variants [11]. This challenge arises due to the different linkage disequilibrium (LD) properties between causal and observed variants for additive and non-additive QTL, where LD of an observed marker tagging a causal variant will manifest at R^2^ for an additive effect, compared to R^4^ for a recessive signal.

This means observed tag variants need to be more closely linked to causal dominance variants to capture the QTL [36,37]. Despite limited prior literature on the use of non-additive GWAS to this end, one noteworthy study suggesting the importance of recessive variants to animal breeding traits was recently reported in the context of male fertility and semen traits in cattle [38]. Here, the researchers identified recessive QTL and candidate causal mutations in several genes including a missense variant in *SPATA16*. That study used imputed genotypes at high density (based on the Illumina BovineHD platform), though it is noteworthy that the study population used was quite small (N=3,736 bulls). It seems likely that the discovery of these QTL was aided in part by the remarkable frequency of the deleterious haplotypes identified in that study, presenting allele frequencies ranging from 9-34% [38].

### Recessive QTL of interest

Although many non-additive signals were identified in this study, we were particularly interested in recessive QTL with large effects, given that these might represent underlying genetic disorders. The five novel recessive QTL on chromosomes 8, 25, 27, and 28 are presented and discussed below.

### Chromosome 8 - DOCK8

Our results present a missense mutation in the *DOCK8* gene as potentially having a deleterious recessive impact on milk yield traits. The QTL appears to operate in a completely recessive manner, with the *DOCK8* variant present at low allele frequencies in each breed (Holstein-Friesian MAF = 0.013, Jersey MAF = 0.059). *DOCK8*, dedicator of cytokinesis 8, is involved with guanine nucleotide exchange factors and influences intracellular signalling networks, and is important in immune responses and lymphocyte regulation in humans and mice [39]. Recessive mutations in *DOCK8* have been associated with hyper Immunoglobulin E syndrome leading to the onset of combined immunodeficiency disease and other health complications [40]. In mice, compromised immune responses are also observed including negative impacts on B cell migration [41], and T cell migration and viability [42,43]. *DOCK8* variants have not previously been associated with cattle performance traits, though if this missense mutation underlies the chromosome 8 QTL, it could be presumed to act through similar negative impacts on the immune system. Under this hypothesis, it is unknown whether the lactation effects are due to mammary immune function or secondary impacts, though given that higher levels of circulating immunoglobulin E and lymphocyte profiling can indicate *DOCK8* deficiency in humans [40,44], future work to sample and profile homozygous animals could be used to definitively establish the causality of the *DOCK8* missense mutation for this QTL.

### Chromosome 25 - IL4R, KIAA0556, ITGAL

The QTL identified on chromosome 25 at 24-27Mbp presented three candidate mutations in genes: *IL4R, KIAA0556*, and *ITGAL. IL4R*, Interleukin 4 receptor, is a transmembrane protein involved in immune responses in humans [45]. *KIAA0556* is associated with microtubule regulation in humans, and knockout mutations in humans and mice have been associated with the neurological disorder, Joubert syndrome [46]. *ITGAL* encodes integrin alpha L chain, and loss of function variants in this gene have been associated with compromised immunity including increased susceptibility to infection to Salmonella in mice [47]. Given that iterative association analysis failed to prioritise one of these variants over the other, it is unknown which of these variants might be responsible for the QTL, and our focus on protein-coding variants as candidates may have also overlooked alternative non-coding or structural mutations as responsible. These variants are nevertheless in moderately strong, though not perfect LD (max. pairwise R^2^= 0.79), so physical genotyping for fine mapping and future functional testing should help to resolve the identity of the gene (or genes) underpinning this QTL.

### Chromosome 25 - LRCH4

Although iterative GWAS did not resolve candidates in the above example, this approach did highlight a second QTL on chromosome 25 represented by a nonsense mutation in the *LRCH4* gene. *LRCH4*, leucine-rich repeats and calponin homology containing protein 4, regulates the signalling of toll-like receptors (TLRs) and has been shown to influence innate immune responses in mice [48]. In that study, researchers showed *LRCH4*-silenced cells presented reduced expression across pro-inflammatory cytokines produced in the TLR4 pathway, most notably that of IL-10 and MCP-1. This suggests a knockout mutation, like that observed here, may have negative impacts on innate immunity in cattle that may drive negative impacts on milk volume, milk fat yield, and milk protein yield.

### Chromosome 27 - SLC25A4

While non-significant at the genome-wide level (c.f. P = 1.65×10^−7^ vs P = 1.30×10^−6^), the chromosome 27 15.5Mbp locus presented a conserved amino acid mutation in *SLC25A4* as the lead associated variant and was therefore of note. This variant demonstrated a complete recessive effect on all three lactation yield traits. The *SLC25A4* gene, solute carrier family 25 member 4, encodes the Adenine nucleotide translocator (Ant1) protein, responsible for the translocation of ATP and ADP between the cytoplasm and mitochondria. In mice, knockouts of *SLC25A4* result in mitochondrial myopathy and cardiomyopathy, and a severe intolerance to exercise [49]. Similarly, in humans, childhood-onset mitochondrial disease and exercise intolerance have been observed for both dominant [50] and recessive mutations [51] in *SLC25A4*. If future association studies confirm the non-significant associations highlighted in the current study, it would be intriguing to examine the phenotypes of homozygous cows further, given the implication that mitochondrial functional deficits and exercise intolerance might underlie these lactation performance impacts.

### Chromosome 28 - RBM34

On first appearance, the strong associations with lactation yield traits near the beginning of chromosome 28 might reasonably be attributed to the *GALNT2* splice site mutation reported and investigated previously [11]. However, upon fitting this mutation as a covariate in our iterative GWAS approach, a secondary peak was still strongly apparent, highlighting a nonsense mutation in the *RBM34* gene as potentially responsible for the effect. The *RBM34* gene encodes an RNA recognition motif protein with an RNA-binding domain. There appears to be little previous research in humans or model organisms on *RBM34*, with limited recent literature probing its involvement in embryonic stem cell differentiation [52]. Here we observed a predicted homozygous knockout of the gene that may influence milk volume, milk protein yield, and milk fat yield in a recessive manner, though its status as a largely uncharacterised RNA-binding protein leaves little room for speculation as to how those effects might manifest. Mechanism aside, identification of two uncorrelated recessive QTL demonstrates the utility of using iterative GWAS approaches, given that ‘peaks’ with compelling causative mutations presented by previous analyses might otherwise go un-investigated. Of further note at this locus, other researchers appear to have observed lactation effects at the 6-10Mb locus previously [53]. However, there appears to be very low LD (R^2^ with RBM34 = 0.04, GALNT2 = 0.02) between the tag variant identified by Raven *et al*. (rs41607517) and the nonsense mutations identified here, suggesting these are likely different effects.

### Previously described additive QTL present partial dominance

We observed several partial dominance QTL closely linked to previously described QTL identified from standard-additive analyses. As described in Supplementary Table 1, we identified dominance components in high LD with variants associated with the genes; *CSF2RB* [33], *MGST1* [16], *DGAT1* [12], *GHR* [13], *AGPAT6* [15], *PLAG1* [54,55], and *PICALM* [34] (and in moderate LD with a *CSN1S1* variant [35]). These partial dominance associations were mostly identified in percentage traits. These observations suggest that many well-known major-effect QTL identified from additive analyses incorporate some level of non-additivity, in agreement with the analyses of milk traits reported by Jiang *et al*. (2017; 2019) [10,32].

## Conclusion

In this study, we have highlighted that different classes of lactation traits (yield compared to composition traits) present differing additive and non-additive genetic architectures. We speculate, that these differences derive from underlying contrasts in the cellular and molecular representation of these traits, where despite comparatively low additive heritabilities, lactation yield effects may better reflect whole-animal energy and fitness status and be a better proxy of a wider range of underlying biological disorders. At a single locus level, we identified five QTL presenting seven candidate causative variants in the *DOCK8, IL4R, KIAA0556, ITGAL, LRCH4, SLC25A4*, and *RBM34* genes, highlighting medium to large effect recessive variants that may provide future opportunity for diagnostic testing and animal improvement.

## Supplementary Figure Legends

**Supplementary Figure 1.**
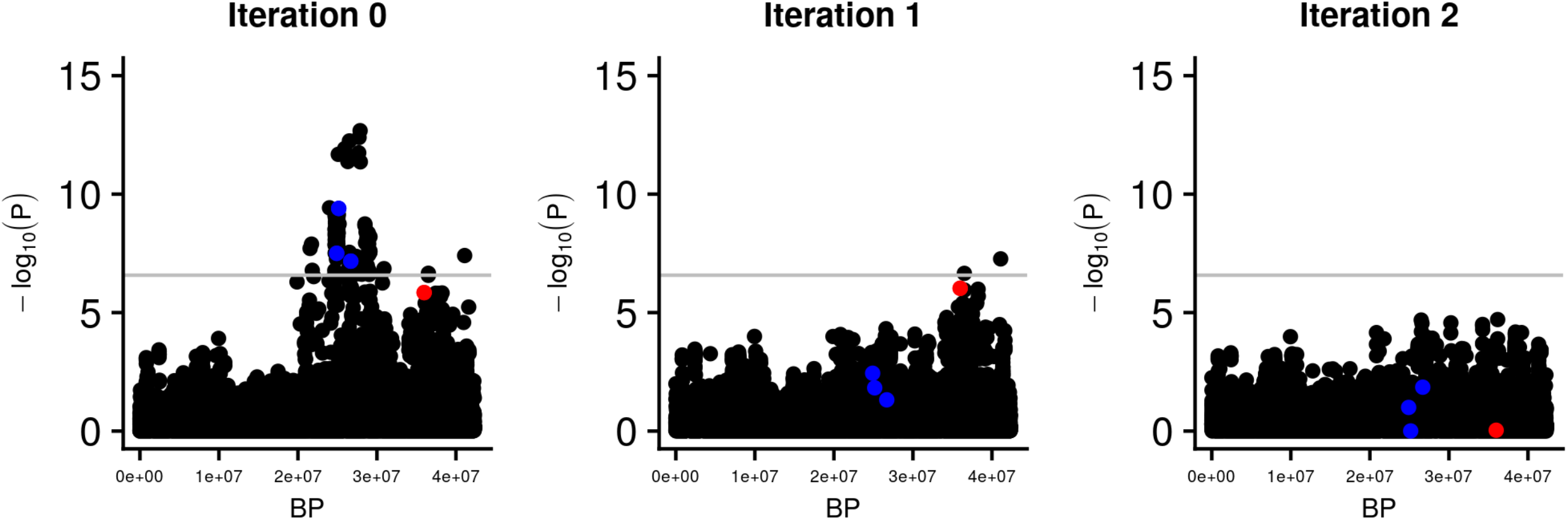
Iterative Manhattan plots for milk-protein yield on chromosome 25. Blue indicates the candidate causal variants in genes; *IL4R, KIAA0556*, and *ITGAL*, and red indicates the candidate causal variant in the *LRCH4* gene. A grey line indicates the false discovery rate of 1×10^−3^, used to account for multiple testing.

**Supplementary Figure 2.**
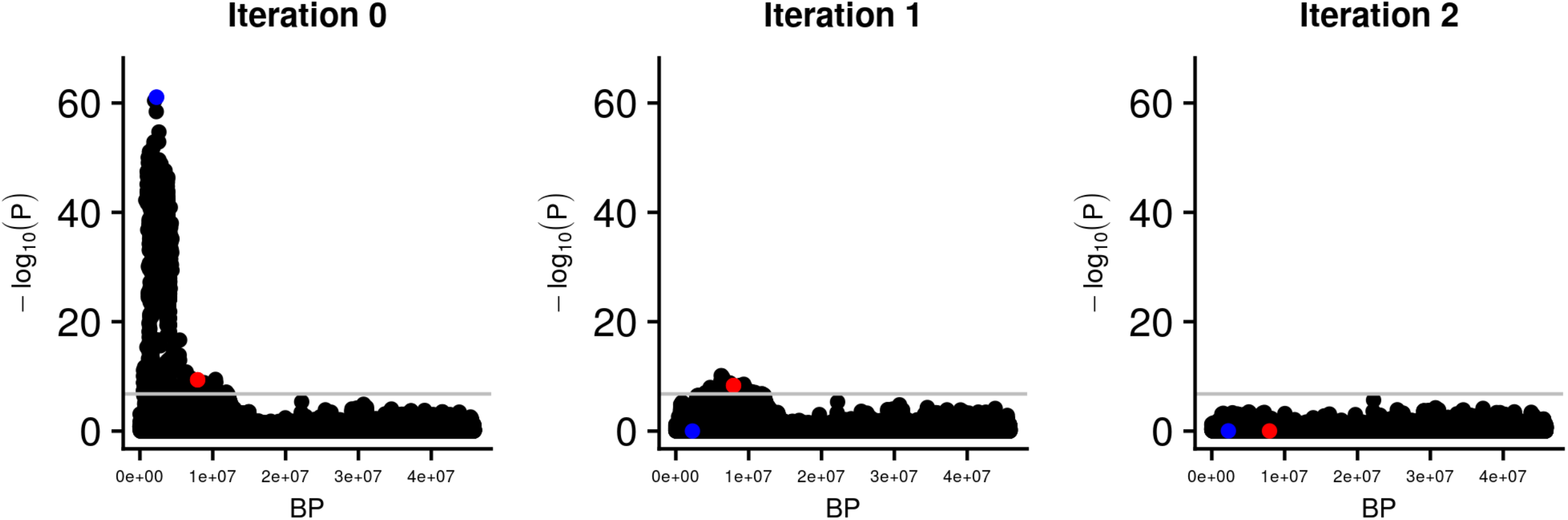
Iterative Manhattan plots for milk-protein yield on chromosome 28. Blue indicates the candidate causal variant in the *GALNT2* gene, and red indicates the candidate causal variant in the *RBM34* gene. A grey line indicates the false discovery rate of 1 ×10^−3^, used to account for multiple testing.

## Declarations

### Availability of data and materials

A subset of whole-genome sequences used for imputation of the genotypes presented in this paper have been deposited in the SRA database [56]. Additional data is available on reasonable request with the permission of Livestock Improvement Corporation, contingent on the execution of an appropriate transfer agreement.

## Acknowledgements

We are grateful for the use of New Zealand eScience Infrastructure (NeSI) high-performance computing facilities as part of this research. We are additionally grateful to the funders of this research.

## Author’s Contributions

EGMR, DJG, and MDL conceived and designed the experiments. EGMR, TL, KT, YW, CSH, BLH, DJG, and MDL performed or assisted statistical analysis. EGMR, TL, KT, YW, CSH, TJJJ, CN, KC, RGS, CC, SRD, BLH, RJS, DJG, and MDL contributed materials or analysis tools. EGMR, DJG, and MDL interpreted results. BLH, RJS, DJG, and MDL were involved in supervising the project. EGMR, DJG, and MDL wrote and revised the manuscript. All authors have read and approved the final manuscript.

## Ethics Declaration

All animal experiments were conducted in strict accordance with the rules and guidelines outlined in the New Zealand Animal Welfare Act 1999. The majority of genotype and phenotype data were generated as part of routine commercial activities outside the scope of that requiring formal committee assessment (as defined by the above guidelines).

